# ModFormer: A unified structure-aware model of transcriptome-wide A-to-I RNA editing

**DOI:** 10.64898/2026.11.23.768611

**Authors:** Cody Slater, Estelle Nassar, Nicholas Dreesen Bonet, Béline Yuan, James Hyojae Lee, Daniel Domović, Lucas Espinosa, Mateusz Garbulowski, Aidan Daly, Sanja Vicković

## Abstract

Adenosine-to-inosine (A-to-I) RNA editing can recode transcripts without permanently altering the genome. ADAR-mediated editing depends on both local sequence and RNA secondary structure, making it difficult to anticipate which sites will be edited or how mutations will change their activity. Moving from observing endogenous A-to-I editing to using it as a programmable molecular tool requires predicting how editing responds to sequence changes. Here, we present ModFormer, a model that integrates RNA sequence representations with predicted secondary structure to predict transcriptome-wide A-to-I editing. Trained on human data, ModFormer predicts sites and rates across substrates and species unseen during training, recovers established sequence and pairing preferences, and anticipates experimentally measured responses to RNA mutations. These predictions prioritize responsive sites in mutational screens and guide an autonomous AI agent toward goal-directed candidate RNA designs. ModFormer provides a framework for predicting how sequence changes reshape RNA editing and can be adapted to new substrates with limited experimental data.

---

Adenosine-to-inosine (A-to-I) editing is an endogenous mechanism of RNA modification that can alter RNA processing, stability, and protein coding potential without changing the underlying genomic DNA. Unlike permanent DNA editing, A-to-I modification is reversible and uses native ADAR enzymes (adenosine deaminases acting on RNA), making it attractive for programmable RNA manipulation [1–7]. However, ADAR editing efficiency depends not only on the nucleotide sequence surrounding a target adenosine, but also on the secondary structure of its RNA substrate, making editing outcomes difficult to predict from the primary sequence alone [8, 9].

ADAR enzymes preferentially act on double-stranded RNA, with editing efficiency shaped by local sequence context, base-pairing interactions, and structural features such as mismatches, bulges, and internal loops [1, 10–12]. These determinants govern editing patterns in endogenous regulation [1, 13] and constrain efforts to engineer targeted RNA correction [4, 14–19]. Because even small sequence changes can alter RNA folding and editing efficiency in ways that are difficult to anticipate, a useful predictive model must go beyond identifying likely edit sites: it must capture their relative activity and anticipate how mutations change editing at a given position. Existing computational approaches have incorporated sequence features, predicted secondary structures, and machine-learning architectures, but their ability to generalize across diverse RNA substrates and experimental contexts remains limited [20, 21].

Recent RNA foundation models offer a path forward by learning sequence and structural dependencies from large-scale pretraining. Models such as RiNALMo learn reusable RNA representations from sequence data, while more recent approaches incorporate structural information directly [22, 23]. However, their training objectives do not capture the biochemical determinants of ADAR substrate recognition or quantify how perturbations influence editing efficiency. Addressing these tasks requires learning from experimentally observed editing outcomes. Large-scale editing catalogs such as REDIportal [24] and direct RNA sequencing measurements of per-site editing rates [25, 26] now provide a foundation for editing-specific predictive models.

Here we introduce ModFormer, a structure-informed RNA language modeling framework that predicts ADAR-mediated A-to-I editing from nucleotide sequence. ModFormer combines representations from pretrained RNA foundation models with predicted secondary structure to learn the determinants of editing-site recognition and efficiency. We evaluate its ability to generalize across species and RNA substrates, test whether its predictions capture the effects of sequence perturbations well enough to guide site selection in mutational libraries, and explore whether an autonomous AI agent guided by ModFormer can search for candidate RNA designs with higher predicted editing. Together, these analyses establish a framework for translating general-purpose RNA representations into editing-specific predictions applicable to programmable RNA sequence design.

## 1 Results

### ModFormer integrates RNA sequence and structure to predict A-to-I editing

We trained ModFormer on human editing data and tested whether it predicts A-to-I sites across species and substrates it had never seen, anticipates the effects of RNA mutations, and guides the computational design of new substrates. ADARs act on double-stranded RNA, including local hairpins and pairing between distant transcript regions [1] (Fig. 1a). We therefore designed ModFormer to represent both the sequence around each adenosine and the region predicted or known to pair with it. To do this, ModFormer combines two interacting representations: a single-sequence track initialized from pretrained language model embeddings (RiNALMo) [22] and a pair track built from base-pairing information (Fig. 1b). ModFormer processes 64-nucleotide tiles within larger sequence windows (up to 1,022 nt), pairing each tile with its predicted duplex partner. Twenty-four Pairformer-style blocks jointly refine these representations, and separate prediction heads estimate editing likelihood and rate (Fig. 1c; Methods).

**Fig. 1.**
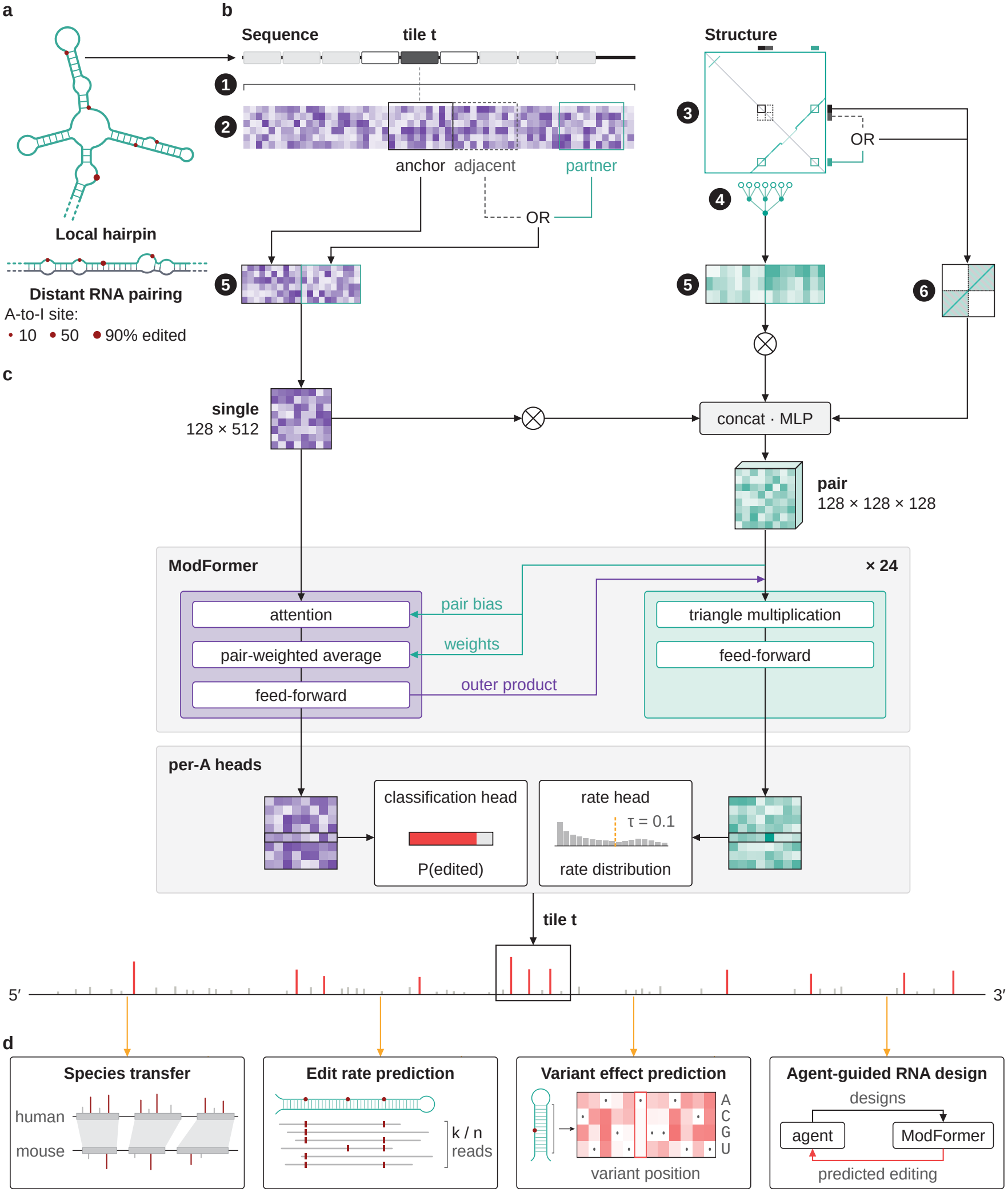
ModFormer integrates RNA sequence and predicted structure to model A-to-I editing. **a**, ADAR substrates formed by local hairpins or distant RNA pairing. Red dots, editing sites; dot size, editing rate. **b**, Input representation: 64-nt anchor tiles are embedded within windows up to 1,022 nt, encoded by frozen RiNALMo and paired with a predicted duplex partner (or adjacent tile). The pair track incorporates predicted base-pairing information. Numbered steps: (1) frozen RiNALMo embedding of the sequence window, (2) tile extraction and projection, (3) base-pairing probability prediction, (4) 1D structure encoding, (5) single and structure representations, (6) pair-representation assembly. **c**, Twenty-four Pairformer-style blocks integrate sequence and pairing representations; separate heads predict editing likelihood and rate (Methods). **d**, Evaluated applications: cross-species transfer, editing-rate prediction, mutation-effect prediction, and agent-guided RNA design. Orange arrows indicate zero-shot or fine-tuned evaluation.

We evaluated ModFormer across species, substrates, and experimental datasets, both zero-shot and after fine-tuning, to test whether its predictions transfer before using them to guide screening and design (Fig. 1d).

### ModFormer predicts editing sites and rates across species and RNA substrates

We first compared ModFormer with six published methods (AdarEdit, InoDREAM, ATTIC, Edit-Predict, PreAIS, and a sequence-neighbor model) on overlap-filtered benchmarks spanning human, invertebrate, and mouse datasets (Extended Data Fig. 1a,b; Methods). ModFormer achieved the highest mean classification AUC (0.92 versus 0.77 for AdarEdit; Extended Data Fig. 1c), including 0.97 in mouse and 0.83 in invertebrates despite training only on human data. On benchmarks drawn from AdarEdit’s and InoDREAM’s own training data, ModFormer matched them closely (AUC 0.93 and 0.94 versus 0.94 and 0.96) without training on either source. ModFormer also ranked measured editing rates more accurately than all comparators in human brain (Spearman *ρ* = 0.24 versus at most 0.08) and mouse (*ρ* = 0.51 versus at most 0.07; Extended Data Fig. 1d).

We next asked whether one epoch of fine-tuning could improve predictions while preserving transcriptome-wide performance. Across seven held-out datasets, joint fine-tuning raised Alu duplex AUC from 0.818 to 0.889 and invertebrate from 0.748 to 0.819, while leaving transcriptome-wide classification AUCs (0.825–0.961 zero-shot) largely unchanged (Fig. 2a; Extended Data Fig. 2a–c; Methods).

**Fig. 2.**
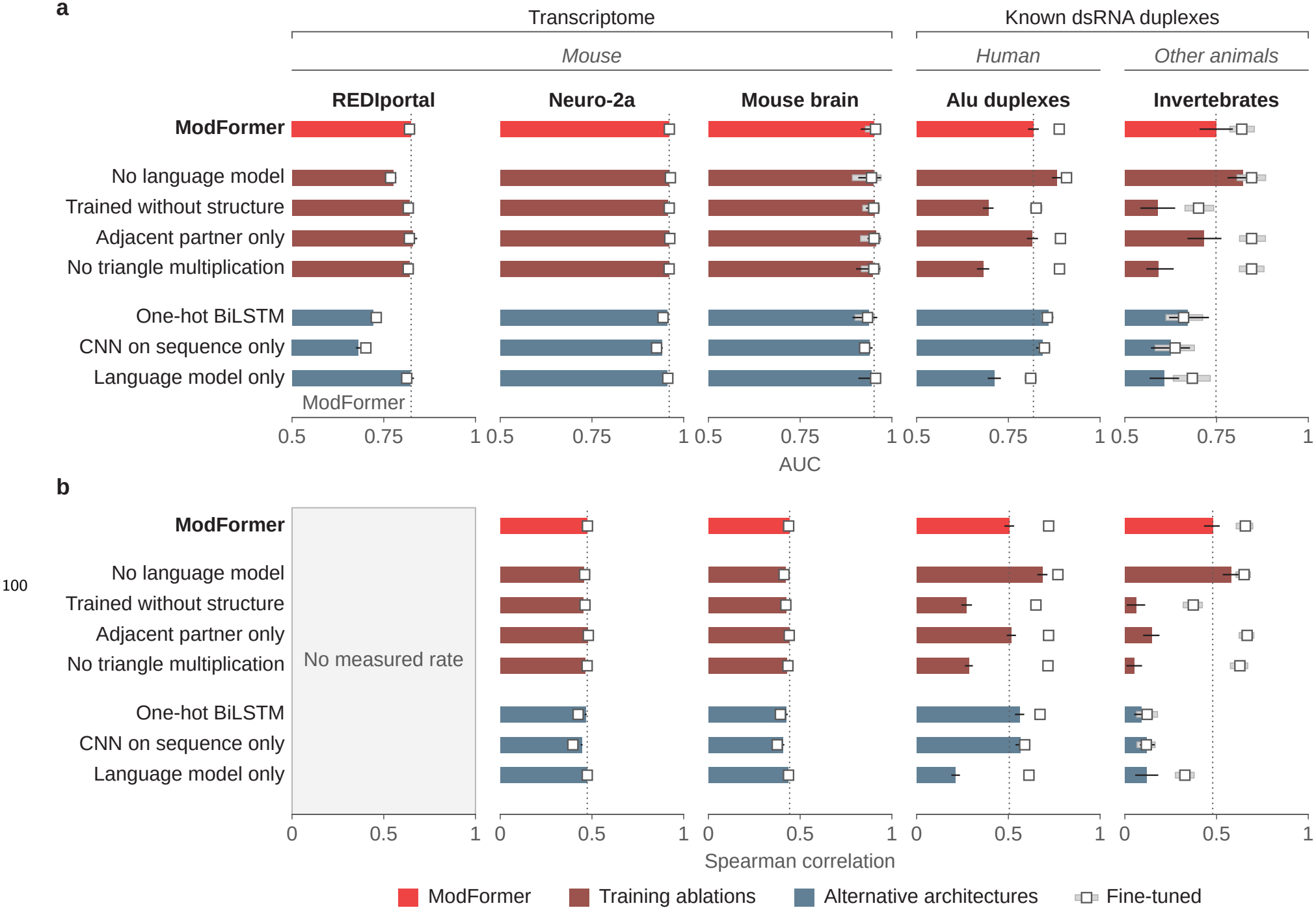
ModFormer predicts editing-site activity across RNA substrates and species. **a**, Editing-site classification AUC across five test sets, zero-shot (bars) and after one epoch of fine-tuning (squares). Dotted lines, full-model zero-shot performance; AUC axes begin at chance (0.5). Thin lines and pale bands, 95% bootstrap intervals. Models grouped by color: ModFormer (coral), training ablations (dark red; no language model, trained without structure, adjacent partner only, no triangle multiplication), and alternative architectures (blue; one-hot BiLSTM, CNN on sequence only, language model only). Test sets span transcriptome-wide (REDIportal, Neuro-2a, mouse brain) and known dsRNA duplexes (human Alu, invertebrate). **b**, Spearman *ρ* between predicted and measured editing rates, including measured zeros; this evaluates site ranking, not absolute rate accuracy. REDIportal lacks measured rates.

Substrate design also requires ranking sites by editing rate. On held-out human and mouse direct-RNA datasets, zero-shot editing-rate correlations ranged from *ρ* = 0.443 to 0.474. For Alu duplexes, *ρ* rose from 0.505 to 0.720 after fine-tuning; for invertebrate duplexes, from 0.479 to 0.657 (Fig. 2b; Extended Data Fig. 2d). The zero-shot model also underestimated absolute editing rates: for human Alu duplexes it predicted a mean of 1.6%, compared with 6.1% measured experimentally; fine-tuning raised the predicted mean to 4.9% (Extended Data Fig. 2e). Fine-tuning therefore improves estimates of a substrate’s expected activity, though agreement between group means does not establish per-site accuracy.

On duplexes, structure drove performance: removing predicted pairing or triangle multiplication dropped zero-shot Alu AUCs to 0.695 and 0.681 (from 0.818), while removing the language-model embeddings left duplex AUCs unchanged or slightly higher (0.882 on Alu). This is consistent with duplex editing being determined primarily by local pairing rather than broader sequence context (Fig. 2a,b). The same pattern held on invertebrate duplexes, where the structure ablations reduced AUC to 0.590 and 0.592 while the no-language-model variant reached 0.821.

By contrast, the language-model embeddings contributed most on transcriptome-wide benchmarks, where sequence-only models scored near chance. The language model was pretrained on single-transcript windows, so some duplex arms overlap a pretraining tile, though in a different structural context. AUCs were comparable with and without such overlap (Extended Data Fig. 2f; Methods), arguing against sequence memorization.

### ModFormer prioritizes active editing sites and recovers ADAR preferences

Alu inverted repeats are among the most common ADAR substrates and span a wide range of editing rates, so they test whether a predictor can reduce the effort of finding active editing sites. We therefore asked how many adenosines must be screened to recover sites with at least 10% measured editing. Across 90 held-out Alu duplexes [21], 2,002 of 11,915 adenosines (17%) met this threshold experimentally. Without additional training, recovering half of these highly edited sites required testing the top 21% of ModFormer-ranked candidates; after fine-tuning, the same recovery required only 14%, approximately one-third fewer candidates (Fig. 3a). When screening the top 17% of model-ranked candidates (equal to the fraction highly edited), ModFormer recovered 41% of active sites before fine-tuning and 57% afterward, compared with 26% and 45% for a sequence-only language-model baseline. We next asked whether the model’s rankings recover known biochemical features. Among mouse REDIportal [24] adenosines ranked in the top 5% for both predicted editing probability and rate (zero-shot, no mouse data seen), the predicted opposing base was cytosine 6.6-fold enriched relative to all evaluated sites, consistent with ADAR’s preference for A–C mismatches [10] (Fig. 3b). Highly ranked sites also showed known sequence preferences: a 2.1-fold enrichment of upstream uridine, a 1.6-fold enrichment of downstream guanosine, and an approximately 20-fold depletion of upstream guanosine [8, 11]. Association alone does not show whether the model responds when those features change. We therefore computationally altered the nucleotide opposite candidate editing sites while retaining the surrounding sequence (Fig. 3c). Introducing an A–C mismatch increased predicted editing, whereas restoring A–U pairing reduced it; nearby substitutions that preserved pairing had little effect. We next tested whether ModFormer predicts the measured effects of deliberate mutations in RNA substrates.

**Fig. 3.**
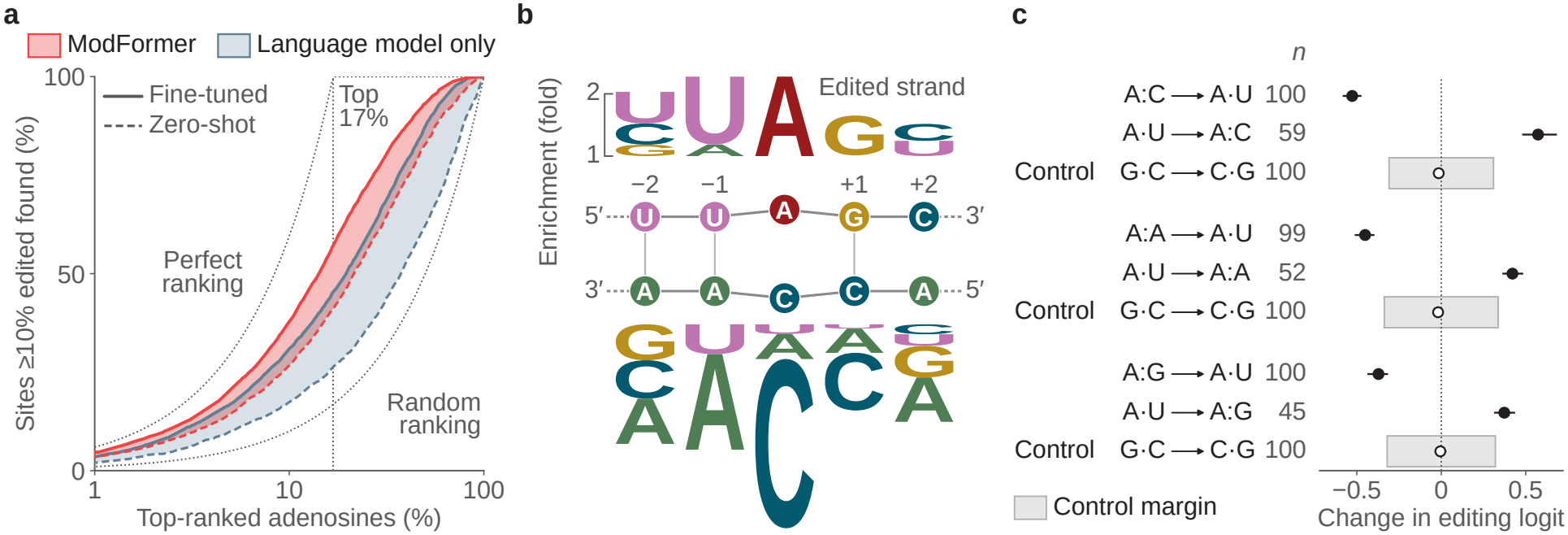
ModFormer prioritizes active editing sites and responds to ADAR-associated sequence and pairing features. **a**, Recovery of highly edited sites (*≥* 10% measured editing; 17% of 11,915 Alu adenosines) versus fraction of candidates screened. Dashed, zero-shot; solid, fine-tuned; gray, perfect and random rankings; dotted line, top 17% of candidates. **b**, Sequence and opposing-base enrichment among 8,775 high-scoring mouse REDIportal adenosines (of 1,293,788 evaluated). Sites rank in the top 5% for both predicted editing likelihood and rate, zero-shot from the human-trained model. Letter height, log_2_ enrichment relative to all sites; top, *±* 2 nt around the adenosine; bottom, predicted opposing base. **c**, Predicted editing log-odds change after computationally creating or removing mismatches opposite the adenosine; nearby pairing-preserving substitutions are controls. A–C introduction increased predicted log-odds by 0.573, A–U restoration reduced them by 0.527, and controls changed them by less than 0.02. Whiskers, 95% family-bootstrap intervals; gray bands, prespecified control margin.

### ModFormer predicts mutation-induced editing changes to guide site selection

We tested whether ModFormer could predict the effects of RNA mutations using a published library of 1,301 synthetic SINE B2 duplexes [27, 28]. Each construct mutated one arm of a reference duplex, and editing was measured at 41 adenosines in the opposing arm to quantify each mutation’s effect (Fig. 4a).

**Fig. 4.**
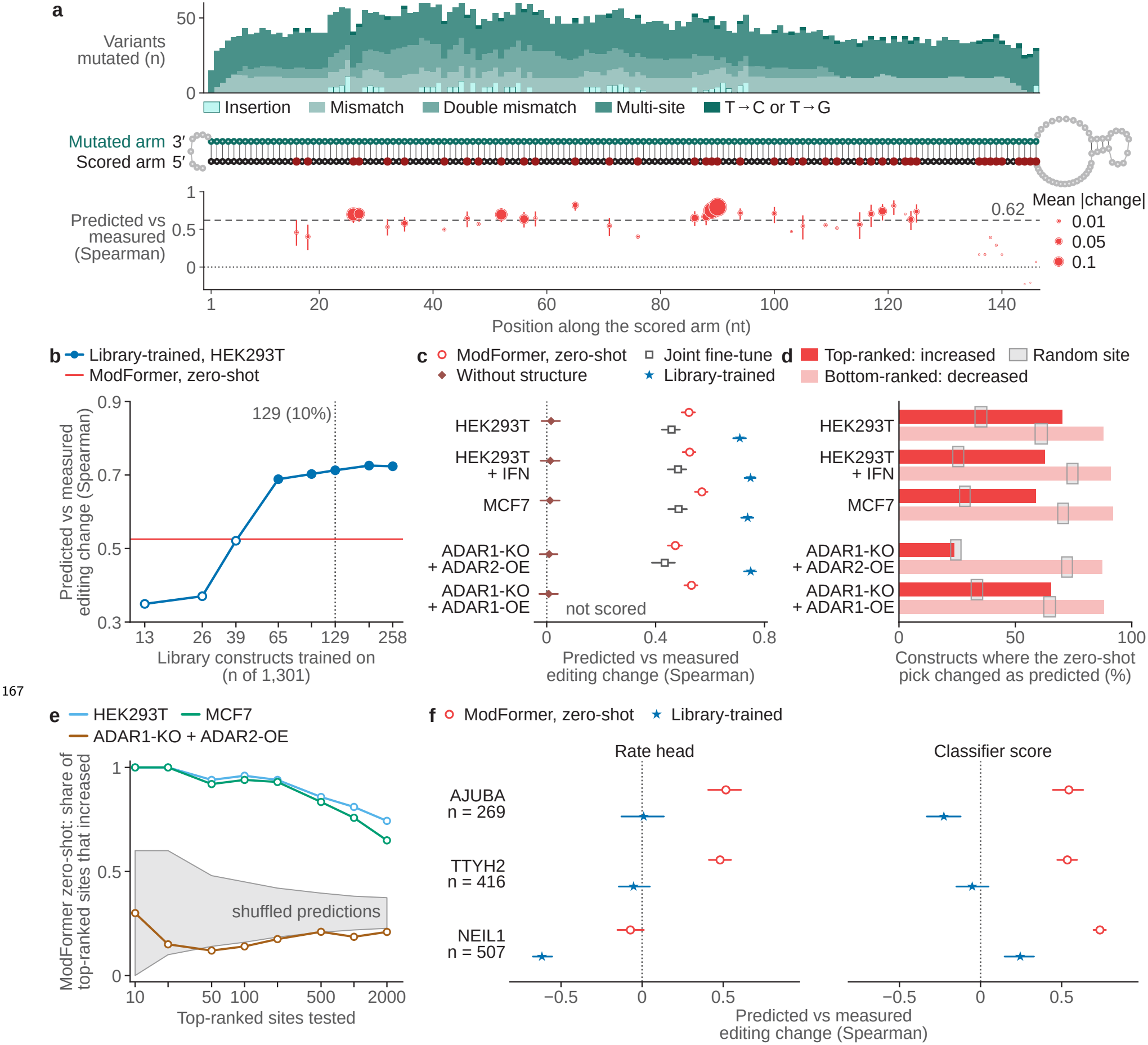
ModFormer predicts mutation effects and prioritizes responsive editing sites. **a**, Synthetic duplex mutations by type and position: teal, mutated arm; black, scored arm; red, edited adenosine. Dot height, zero-shot Spearman *ρ* between predicted and measured editing changes in HEK293T; area, mean absolute change; whiskers, 95% bootstrap intervals. Dashed line, pooled *ρ* (0.62 across 4,559 mutation-proximal adenosines). **b**, Editing-change *ρ* versus library-training budget; dotted line, 129 designs (10%). Filled points, significant improvement over zero-shot (95% bootstrap); open points, not significant. **c**, Editing-change *ρ* across four models and five cellular conditions, with ADAR1-knockout conditions distinguished. Whiskers, 95% intervals. **d**, Fraction of constructs whose predicted highest-gain or highest-loss site changed editing in the expected direction (zero-shot); gray, 2,000 random selections per construct. **e**, Fraction of ranked sites with measured editing gains versus number tested; curves, ModFormer zero-shot, one per condition; gray, 2,000 shuffled rankings. **f**, Spearman *ρ* between predicted and measured editing changes for endogenous AJUBA (269 variants), TTYH2 (416), and NEIL1 (507) substrates [29]. Red circles, zero-shot ModFormer; blue stars, SINE B2 library-trained predictor; each scored with both editing-rate and classification predictions.

Without training on this library (zero-shot), ModFormer predicted editing changes with a correlation of *ρ* = 0.62 across 4,559 mutation-proximal adenosines in HEK293T cells (95% confidence interval 0.60–0.65; Fig. 4a; Methods). We then asked how few library designs a model would need to outperform the zero-shot baseline. Training a model from scratch on a random subset of the library improved performance with as few as 65 duplexes (2,665 A’s) and had largely plateaued by 129 (5,289 A’s), approximately 10% of the full library (Fig. 4b). With 129 duplexes for training and 1,172 held out, the correlation rose to *ρ* = 0.709 (0.685–0.732; Fig. 4c). Adding those same 129 designs to the broader joint fine-tuning (Extended Data Fig. 2b) had the opposite effect, reducing the correlation to *ρ* = 0.459 (0.423–0.489) below the zero-shot baseline. A small library-specific training set thus outperformed the zero-shot model, but the same data diluted into broader training hurt performance.

A high correlation across thousands of sites does not, by itself, tell an experimenter which site in a given construct to target. We therefore asked whether ModFormer could identify the most responsive adenosine in each design. In HEK293T cells, the top predicted site gained editing in 70.3% of constructs, versus 35.3% for a random site; the bottom predicted site lost editing in 87.9%, versus 61.0% for random (Fig. 4d). Similar patterns held in interferon-treated HEK293T and MCF7 cells. The selected sites lay 20–40 nucleotides from the mutation’s pairing partner (Extended Data Fig. 3a), consistent with the ADAR binding footprint [12], and outperformed random sites at the same distances (Extended Data Fig. 3c). Ranking multiple sites per construct and pooling across designs enriched for measured editing gains (Fig. 4e; Extended Data Fig. 3d,e,f).

Mammals express two catalytically active editing enzymes, ADAR1 and ADAR2, with different substrate preferences [9]. The same SINE B2 library was independently measured in ADAR1-knockout cells overexpressing either enzyme [28], letting us ask how ModFormer’s predictions fare when the enzymatic context changes. Library-trained predictions still correlated with measured editing changes under ADAR2 overexpression (*ρ* = 0.749; Fig. 4c), but site selection did not transfer: zero-shot ModFormer identified a gaining site in only 23.7% of constructs, comparable to 24.5% for random (Fig. 4d; Extended Data Fig. 3b). Under ADAR1 overexpression, the top predicted site gained editing in 65.3% of constructs, compared with 33.5% for random. Predicting mutation effects across a library and identifying which site to engineer in a given RNA are therefore distinct challenges, with site selection particularly sensitive to the ADAR context.

We tested generalization to endogenous substrates using published mutational scans of AJUBA (269 variants), TTYH2 (416), and NEIL1 (507) [29] (Fig. 4f). Without training on these libraries, ModFormer’s classification scores correlated with measured editing changes across all three substrates (Spearman *ρ* = 0.54, 0.54, and 0.74). Editing-rate predictions also correlated with measured changes in AJUBA (*ρ* = 0.52) and TTYH2 (*ρ* = 0.48) but not NEIL1 (*ρ* = *−*0.07), where the model underestimated editing rates. By contrast, the model trained on the synthetic SINE B2 library showed near-zero or negative correlations on all three endogenous substrates (Fig. 4f). The zero-shot model thus captures transferable sequence–structure information, whereas library-specific adaptation improves performance on its training substrate at the cost of generalization.

### ModFormer enables agent-guided computational RNA design

We gave a general-purpose AI agent access to the zero-shot model and asked it to redesign the partner arm of a synthetic SINE B2 duplex to increase editing at a specified adenosine. The target sat in a 5’-GAG-3’ context that disfavors ADAR editing [11], and its neighboring nucleotides remained fixed, so the agent had to explore cross-strand changes and overall duplex structure. It evaluated 1,000 distinct designs, receiving only ModFormer’s predicted editing score and whether the target adenosine was paired or unpaired (Fig. 5a; Methods).

**Fig. 5.**
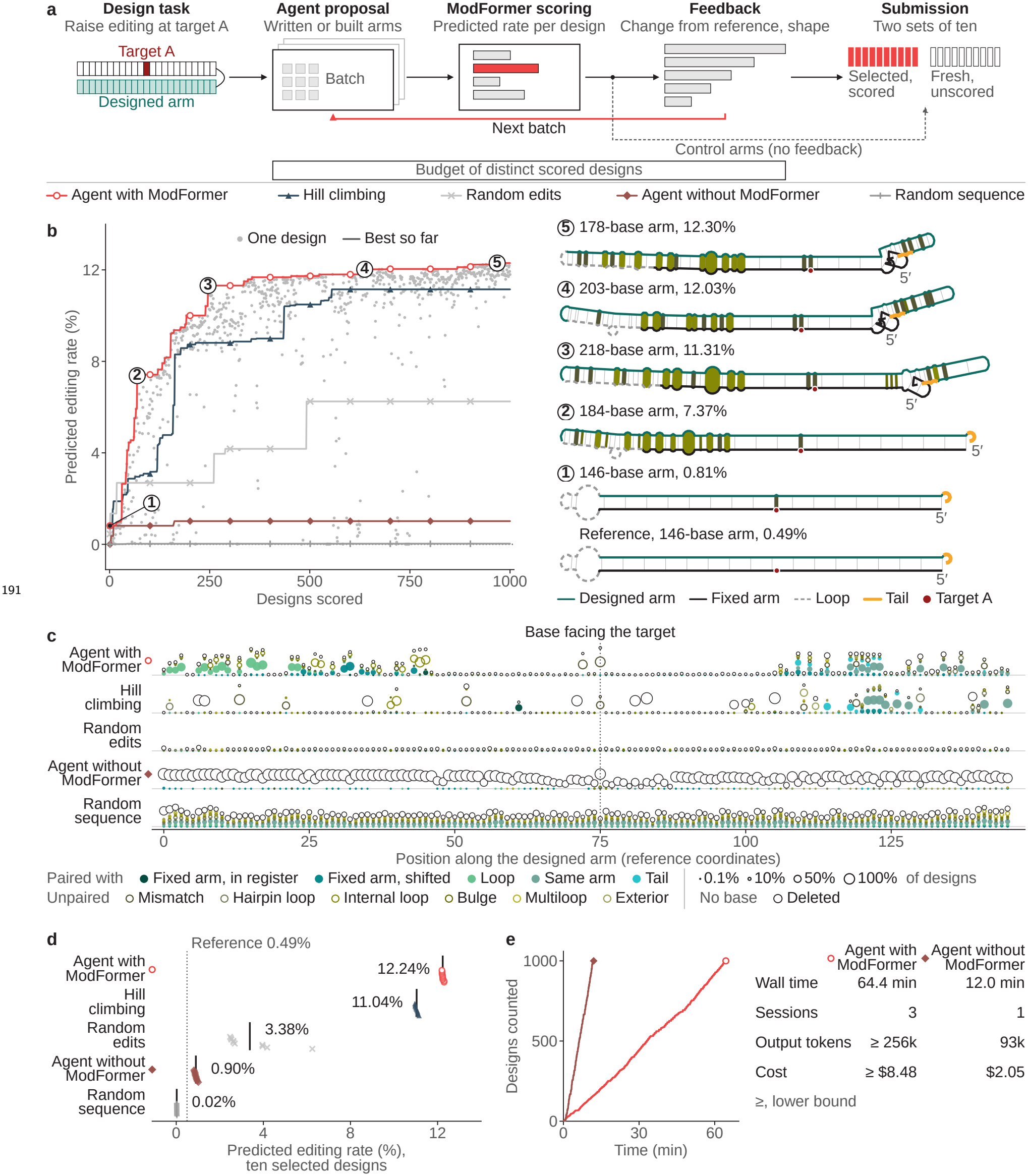
ModFormer-guided computational search prioritizes changes opposite the target adenosine. **a**, Five design strategies with a budget of 1,000 distinct designs each. Guided agent (red open circle) and hill climbing (slate triangle) receive ModFormer scores during search; blind agent (muted red diamond), random edits (*×* ), and random sequences (+) are scored afterward. **b**, Predicted target editing (left) from the reference (0.49%) to the best agent design (12.30%). Examples (right), designs 1, 69, 246, 637, and 968; red dot, target adenosine, paired in the reference minimum-free-energy fold and unpaired in the examples. Folds share a scale; mismatches, internal loops, and bulges are filled in the colors of **c. c**, Sequence substitutions and predicted pairing by strategy; rows are offset by strategy. The position opposite the target was substituted in 924/1,000 guided-agent, 559/1,000 hill-climbing, and 46/1,000 random-edit designs. Open circles, deletions; dot area, fraction of designs. Counts describe sequence outcomes, not agent intent. **d**, Mean predicted editing of each strategy’s ten selected designs: guided agent, 12.24%; hill climbing, 11.04%; random edits, 3.38%; blind agent, 0.90%; random sequences, 0.02%. Dotted line, reference. Both guided methods optimize the evaluation score; no designs were experimentally tested. **e**, Guided versus blind agent: 64.4 versus 12.0 min, *≥*256,080 versus 93,283 output tokens, and *≥*$8.48 versus $2.05.

The agent placed a mismatch opposite the target adenosine in its very first design. Early designs carried large internal loops far from the target and scored near zero. By design 31, the agent had eliminated those loops and scores began to rise. By design 192 (predicted editing rate at 9.5%), the search had converged on a recurring template: a 1*×*1 mismatch at the target, a second mismatch two nucleotides away and an internal loop *∼*28 nt downstream, leaving the target unpaired in the minimum-free-energy fold (Fig. 5b). Of the remaining 808 designs, 94% shared this template; the best reached 12.30%, up from 0.49% for the reference duplex.

We compared this guided agent against four baselines, all given the same 1,000-design budget: score-guided hill climbing, which also receives ModFormer feedback; a blind agent that designs without scores; random single-nucleotide edits; and random sequences (Fig. 5a). The guided agent substituted the position opposite the target 92.4% of the time, compared with 55.9% for hill climbing and 4.6% for random edits (Fig. 5c). Its ten selected designs averaged 12.24% predicted editing, exceeding hill climbing (11.04%), random edits (3.38%), the blind agent (0.90%), and random sequences (0.02%; Fig. 5d). The agent needed no manually specified mutations, converged on a productive template within about 200 designs, and finished in 64.4 minutes at a cost of at least $8.48 (Fig. 5e). No designs were experimentally tested.

## 2 Discussion

ModFormer demonstrates that a single sequence–structure representation, trained on human editing data, can support three complementary objectives: accurate prediction of endogenous editing sites, rapid adaptation to specialized substrates, and computational prioritization of candidate RNA designs. These capabilities arise from jointly learning sequence and structural features relevant to ADAR substrate recognition, rather than relying on sequence preferences or structural rules alone.

The results reveal a context-dependent relationship between sequence and structure in predicting editing. Language-model embeddings contributed most to transcriptome-wide site identification, consistent with the dominant role of local sequence context in determining whether an adenosine is edited. Removing predicted secondary structure had a larger effect on characterized duplexes, where base-pairing geometry directly shapes ADAR access and efficiency. Cross-species generalization from human training data to mouse and invertebrate RNAs suggests that the features governing ADAR recognition are at least partially conserved. Within a single substrate system, library-specific training improved predictions on held-out variants, reducing the experiments needed to identify responsive sites. However, the zero-shot model outperformed the library-trained model on unrelated endogenous ADAR1 substrates, indicating that library-specific training can overfit to library-specific patterns at the expense of the broader recognition grammar. This tradeoff between substrate-specific precision and general applicability suggests that the choice of model should match the experimental context. The agent-guided design experiments illustrate a broader opportunity: a predictive model that captures sequence–structure relationships can serve as the objective function for autonomous sequence optimization. Given only predicted editing scores and pairing state, the agent independently converged on a structural template consistent with known ADAR preferences, including a mismatch opposite the target adenosine and a nearby internal loop, without being told what features to explore. This is qualitatively different from screening a fixed library or applying hill climbing on individual positions: the agent reasons about how coordinated sequence changes reshape RNA structure and, through it, predicted editing. Although the resulting designs are computational candidates requiring experimental validation, they draw on substrate-recognition rules learned from a large corpus of measured editing data to define starting points that manual enumeration alone could not reach. This paradigm extends beyond the SINE B2 substrate used here. Any editing target for which ModFormer provides predictions, whether endogenous or engineered, human or cross-species, can in principle be explored through the same goal-directed search, with the model’s generalization determining how far the designs can extrapolate from training data.

Several limitations should be noted. ModFormer was trained exclusively on human editing data and does not model cell-type or condition-dependent variation in ADAR expression or interacting RNAs. Editing rate predictions, while correlated with measured rates, remain approximate. The current model uses predicted secondary structure as a fixed input rather than learning structure jointly, which may limit its sensitivity to subtle structural perturbations. Coupling the agent with experimental feedback, testing a subset of proposed designs and retraining on the results, could close the loop between computational search and measured editing outcomes.

These results establish a framework for translating general-purpose RNA representations into editing-specific predictions. By capturing the relationship between sequence, structure, and editing outcome, ModFormer provides a starting point for predicting how sequence changes will reshape editing, a step toward the autonomous, goal-directed design of programmable RNA editing substrates.

## Online Methods

### Datasets

Datasets for pretraining, joint fine-tuning, library-specific adaptation, and external evaluation are summarized in Supplementary Table 1 and Extended Data Fig. 2a.

### Pretraining

Pretraining sources: human REDIportal binary editing annotations [24] and HEK293FT direct RNA sequencing data, sampled at a 5:1 ratio. REDIportal labels reflect catalog membership with no read-support threshold. Direct-RNA classification thresholds: *≥*20 valid reads to be measured, rate *≥*0.10 to be positive. The rate head trained on all sites with *≥*1 covering read, including measured zeros, because the binomial mixture loss weights each observation by its coverage.

### Fine-tuning

Joint fine-tuning added mouse REDIportal, Neuro-2a, and mouse-brain direct RNA, and characterized human Alu and invertebrate duplexes [21]. Library-specific training on 129 synthetic SINE B2 constructs was evaluated separately (Designed-library screen, below). Natural-substrate mutational scans were not used for training.

### Data partitions and overlap

All datasets: 80:10:10 train/validation/test split, separated by gene. The Alu duplex test set was stratified post hoc by pretraining sequence overlap: a duplex was considered overlapping when either arm shared a 128-nt window with a labeled pretraining tile (Extended Data Fig. 2f). Published Alu and invertebrate duplex measurements were re-partitioned for these analyses.

### External cohorts

Synthetic duplex mutation libraries [27, 28], endogenous ADAR1 substrate scans [29], and published-model benchmark datasets were filtered under the inclusion rules in Comparison with published models. No natural-substrate scans were used for training. Source accessions, cohort sizes and dataset roles are in Supplementary Table 1.

### Direct RNA sequencing

Rate measurements for HEK293FT, Neuro-2a and mouse brain come from Oxford Nanopore direct RNA sequencing. HEK293FT cells (Thermo Fisher, R70007) and Neuro-2a cells (ATCC, CCL-131) were cultured in DMEM with 10% fetal bovine serum at 37 °C and 5% CO_2_. Brain tissue came from one male C57BL/6J mouse at 2 months of age, euthanized by cervical dislocation; the brain was flash-frozen on dry ice. All animal procedures were performed in accordance with protocols approved by the Columbia University Institutional Animal Care and Use Committee. Total RNA was isolated with TRIzol (Invitrogen, 15596026) according to the manufacturer’s protocol. Libraries were prepared from 1 µg of total RNA with the Oxford Nanopore Direct RNA Sequencing Kit (SQK-RNA004), following the manufacturer’s protocol for reverse transcription and adapter ligation, and sequenced on PromethION RNA flow cells (FLO-PRO004RA). Total RNA was sequenced on five sequencing runs from two replicates for HEK293FT, six sequencing runs from three replicates for Neuro-2a, and three sequencing runs for mouse brain. All reads were basecalled with Dorado 1.0.0 using the rna004_130bps_sup@v5.2.0 model, aligned to the genome (GRCh38 or mm39) with splice-aware minimap2 2.31 (`-ax splice -k14 -uf –MD –secondary=no -y`), and summarized per position with modkit 0.6.4 pileup, filtering to the inosine modification channel (code 17596). Each adenosine carries separate inosine, m6A and 2’-O-methyl-A calls; reads called m6A or 2’-O-methyl-A remain in the valid-read count but are not counted as edits. The adenosine pass threshold was set at the 10th percentile of inosine modification probabilities per sample (modkit sample-probs; 0.76 for HEK293FT, 0.67 for Neuro-2a, 0.77 for brain). A valid read is one that passes modkit’s base-quality and threshold filters (bedMethyl *N*_valid_). A site is measured at *≥*20 valid reads and positive at a rate *≥*0.10.

### Structure prediction

Folds were computed with ViennaRNA [30] (version 2.7.2; 2.7.1 for the HEK293FT precompute) over the same window and RNA strand as the sequence track. Settings: 37 °C, 0.15 M salt, dangles 2. Outputs: the minimum-free-energy structure and the partition-function base-pair probabilities. All structures are thermodynamic predictions from single-strand folds; no experimental probing data were used.

### Architecture

Each example is a 128-nt crop made of two 64-nt tiles. The first is the anchor, whose adenosines are all scored exactly once per epoch. The second is the partner, which provides structural context but is never scored. The partner is the 64-nt window centered on the anchor’s dominant predicted helix. To find it, anchor positions whose maximum base-pair probability exceeds 0.5 with a partner outside the anchor are grouped by register; the largest group defines the partner window. If no anchor position has a confident off-anchor partner, the adjacent 64 nt are used. The window that is folded and embedded around the crop is 1,022 nt for the transcriptome sets, 704 nt for the Alu duplex sets, 384 nt for the reporter-library constructs and 128 nt for the Liu substrates (Fig. 1b).

#### Single track

Sequence enters as embeddings from RiNALMo-giga [22], a frozen RNA language model with 650 M parameters, 33 layers, and 1,280 hidden dimensions, loaded from the released RiNALMo-giga weights via `multimolecule` 0.2.0. Embeddings are precomputed over 1,022-nt windows at a stride of 128, giving a 1,280-dimensional vector per nucleotide, which a learned projection (658,432 parameters, trained with the model) maps to 512 (Fig. 1b,c). **Pair track**. A one-dimensional structure encoder (1,237,872 parameters, trained with the model) reads thirteen per-position summaries of the base-pair probabilities and a six-class loop-type label. It never sees the base-pair matrix or the sequence. Its 256 channels per position are indexed to the 128 anchor-and-partner positions. Two outer-product branches of 256 channels each, one from the single track and one from the structure encoder, are concatenated with 48 pairwise channels (nine pairwise relations, including the base-pair probability). A two-layer cell-wise multilayer perceptron reduces the 560 channels to a pair width of 128, giving a 128 *×* 128 *×* 128 pair tensor.

#### Trunk

The two tracks enter 24 Pairformer-style blocks [31] (Fig. 1c). Single-track width: 512. Pair-track width: 128. Attention heads: 16. Feed-forward width: 4*×* the track width. Dropout: 0.1. Each block runs pair-biased single attention, a pair-to-single update, a single feed-forward layer, a single-to-pair write, outgoing and incoming triangle multiplication, and a pair feed-forward layer. The blocks depart from the published Pairformer in four ways: (1) triangle multiplication in both directions, no triangle attention; (2) SwiGLU transitions at two-thirds of the track width; (3) a zero-initialized bilinear single-to-pair write of rank 32 and a pair-to-single update where the pair track owns the softmax and the single track supplies the values; (4) RMSNorm with weights in fp32. The released checkpoint has 116,633,234 trainable parameters, not counting the frozen language model.

#### Heads

Two per-adenosine heads read three inputs: the single vector at the scored adenosine, an attention-pooled row of the pair representation, and the pair diagonal (Fig. 1c). This gives a direct pair-to-output path alongside the single track. A classification head gives an edit detection score. A rate head gives a distribution over editing rate in 17 bins, and the point estimate is the expected value of those bins. The threshold *τ* = 0.1 defines the positive label and the boundary between the last finite bin and the zero atom. It does not enter the point estimate.

### Training

#### Pretraining

Training configurations, optimizer-step counts, architecture dimensions, and finetuning settings are summarized in Supplementary Table 2. The total loss combines classification, within-tile ranking, and measured-rate terms with equal weight,

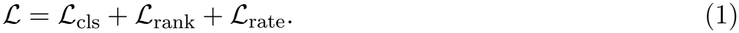

The classification head is a separate multilayer perceptron trained with cross-entropy, *L*_cls_, against catalog membership, that is whether REDIportal lists the site. The ranking term runs on the same channel. For each edited adenosine *e* in a tile it draws five less-edited adenosines *u* from the same tile at random and applies a hinge with margin *m* = 0.1 to the classification scores *s*,

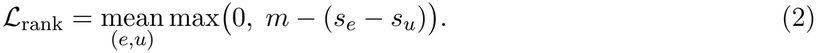

The rate head gives probabilities *q*_*b*_ over 17 rate bins. For a site with *k* edited reads out of *n* covering reads, the rate loss is the negative log-likelihood of a binomial mixture over the bins,

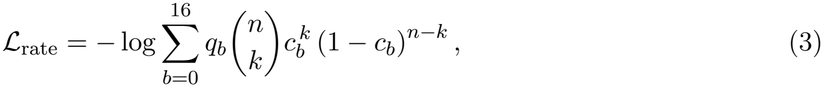

The 17 bins comprise a zero atom and 16 finite bins whose edges form two geometric series meeting at 0.10 (step ratio 1.631 below, 1.334 above); *c*_*b*_ is the geometric mean of each bin’s edges (*c*_0_ = 0). The loss is evaluated on all measured sites with *n ≥* 1, including measured zeros.

#### Joint fine-tune

The joint fine-tune has the same architecture settings as the base model and adds a learned context vector, the sum of learned assay, organism, and tissue embeddings, to the inputs of both heads. It trains all weights for one epoch with an L2-SP penalty [32] that pulls each inherited parameter *θ*_*i*_ toward its pretrained value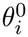,

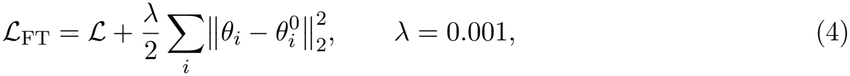

where the sum runs over trainable inherited parameters, including biases and normalization gains, and excludes newly added parameters such as the context embeddings. The fine-tune draws from the fine-tuning and pretraining sources at the shares shown in Extended Data Fig. 2b.

### Evaluation

Detection metric: area under the receiver operating characteristic curve (AUC), with average precision reported alongside it. The scoring head depends on the label type: where labels are measured editing rates, the detection score is the rate head’s probability that a site exceeds the rate cutoff; otherwise it is the classifier probability. In Fig. 2a, the rate head scores Neuro-2a and mouse brain; the classifier scores the other three sets. The same rule applies to published-model comparisons (Extended Data Fig. 1b–d).

Rate metric: Spearman correlation between predicted and measured rates over positions with *≥*1 read (995,381 adenosines in HEK293FT), including measured zeros. This assesses ordering across sites, not absolute-rate accuracy. The bin-averaged comparison (Extended Data Fig. 2e) shows calibration on the rate scale.

### Site-screening recovery

The recovery analysis (Fig. 3a) ranks all 11,915 adenosines across 90 held-out Alu duplexes by the rate head’s expected editing rate, pooled across duplexes. A site is highly edited if its measured rate *≥*0.10 (2,002 of 11,915, 16.8%). The recovery curve plots the fraction of highly edited sites recovered as a function of the fraction of candidates screened. Reference curves are analytic: the random ranking follows *y* = *x*; the perfect ranking follows *y* = min(1, *x/*prevalence).

### Ablations

Four ablations were trained with the same corpus, schedule and seed as the released checkpoint: (i) shuffled fold (permuted upper triangle of the base-pair matrix), (ii) no pair track, (iii) adjacent window in place of the structural partner, (iv) no triangle multiplication. A fifth, the no-language-model ablation, replaces RiNALMo embeddings with a learned nucleotide embedding. All were evaluated on the held-out test sets (Fig. 2a). Three alternative architectures used the same corpus and schedule: a one-hot BiLSTM (2 layers, hidden 256 per direction, 2.4 M parameters), a parallel-kernel CNN on sequence only (2.6 M parameters), and a language-model-only transformer (2 layers, 256-dimensional, no structure input, 2.1 M parameters).

### Sequence and opposing-base enrichment

High-scoring sites (Fig. 3b): the 8,775 mouse REDIportal adenosines ranking in the top 5% for both predicted editing likelihood and predicted editing rate (zero-shot), out of 1,293,788 evaluated.

For each position within *±*2 nt of the candidate adenosine, enrichment is the log_2_ ratio of the base frequency among selected sites to its frequency among all evaluated sites. The same all-site background was used for the predicted opposing base.

### Comparison with published models

Comparators: AdarEdit [21], InoDREAM [33], ATTIC [34], EditPredict [35], PreAIS [36], and a sequence-neighbor rule [11], all evaluated with released weights and code where available. To avoid training-data overlap, each benchmark retained only records absent from the training data of every evaluated model (for ModFormer, sites outside any pretraining input window). Source checkpoints, input lengths, filtering counts, and benchmark-specific label definitions are in Supplementary Table 3. We evaluated the published models on the authors’ Alu and InoDREAM test records, the invertebrate duplex benchmark and a sampled set of mouse Neuro-2a and brain sites. These are distinct evaluation cohorts from those used in Fig. 2a, with different biological sources, class labels, and sampling (cohort definitions in Supplementary Table 3). To ensure comparable inputs in the Alu benchmark, single-sequence models were evaluated only at sites for which their required centered sequence windows could be extracted after removing the duplex linker. AdarEdit was supplied its required duplex or folded window as appropriate.

Editing-rate comparisons (Extended Data Fig. 1d) used tissue-specific rates from InoDREAM Supplementary Dataset S2 [33]: HEK293T (2,461 sites) and human brain (775 sites), with replicate-summed coverage *≥*100. Uncertainty: 1,000 percentile-bootstrap resamples, clustered by duplex, species, gene, or record as specified in Supplementary Table 3; between-model comparisons used paired resamples.

### Designed-library screen

#### Library design and editing measurements

The library contains 1,301 designs, each mutating one arm of a reference duplex and measured on the other arm, in HEK293T cells, HEK293T cells with interferon, and MCF7 cells [27] (BioProject PRJNA649938), and in ADAR1-knockout cells with ADAR2 or ADAR1 overexpression [28] (Bio-Project PRJNA943413); construction and measurement are as described in those studies. Editing change is each variant’s measured rate minus the reference construct’s rate at the matched position, without normalization.

#### Library-trained models and site prioritization

Library-trained models were fitted on 129 constructs and evaluated on the 1,172 held out, using a single family-stratified split. The budget curve (Fig. 4b) retrains at each budget and scores on the same held-out constructs; a filled marker indicates a budget whose paired interval against the released checkpoint lies above zero. Each construct’s pick is taken over the 41 adenosines on the conserved arm, the same positions in every construct; adenosines on the mutated arm are scored but are not candidates for the pick. Sites are ranked by signed predicted change in editing rate; the top-ranked site is scored for an increase and the bottom-ranked for a decrease. Top-pick and random-pick comparisons (Fig. 4c–e, Extended Data Fig. 3b,d,e) use two different nulls and are not interchangeable: one-pick bands come from 2,000 within-construct random picks, and shuffled-prediction bands from 2,000 shuffles of the predictions.

#### Mutation distance and distance-only baseline

Editing is measured on arm 1 and mutations lie on arm 2, so no scored site is within 46 nt of a mutation in sequence. Because the duplex pairs arm positions summing to 336, a mutation at arm-2 position *q* pairs with arm-1 position 336 *− q*. Distances are measured along arm 1 to the nearest such position.

#### Analysis cohorts and plotting conventions

The evaluated sets are 7,734 sites in 1,034 constructs for Fig. 4b and 8,636 sites in 1,163 held-out constructs for Fig. 4c, whose whiskers are 95% intervals over 1,000 construct resamples; the dashed line in Fig. 4a, 0.62, is pooled over 4,559 site-design pairs, and 1,301 to 1,302 constructs carry a pick in each condition in Fig. 4d. In Fig. 4a, insertions are counted at the base 5^*′*^ of the junction they sit in, positions are 1-based, each dot reads the designs whose nearest mutation lies within 3 nt of that adenosine, and a dot is drawn where the mean absolute measured change exceeds 0.01. All 1,301 designs are read, including the 129 that trained the library-trained model. The reference and agent-design folds in Fig. 5b are minimum-free-energy folds from ViennaRNA 2.7.2 at 37 °C and 0.15 M salt.

#### External mutational scans

The three natural substrates (AJUBA, TTYH2 and NEIL1; Fig. 4f) are deep mutational scans from Liu et al. [29] (GEO GSE138860); their construction is described there. Each substrate was scored with both the zero-shot pretrained checkpoint and the SINE B2 library-trained model, using both the classification and rate prediction heads (Fig. 4f).

#### Comparisons across cellular and ADAR contexts

The five cellular conditions come from two studies of the same design pool, not from one experiment: 1,301 constructs share identifiers and arm sequences with HEK293T. Across the HEK293T/MCF7 and knockout groups, study and cell background differ alongside enzyme context. The two ADAR1-knockout overexpression conditions share a knockout background and permit a more direct contrast between ADAR1 and ADAR2 contexts, but the available comparisons do not establish equivalent enzyme expression or activity. Fig. 4d reports zero-shot top-site selection across all five conditions. By contrast, Extended Data Fig. 3b shows library-trained predictions in the first four conditions and zero-shot ModFormer in the ADAR1-overexpression condition.

#### Target-pairing perturbations

We compared predicted editing log-odds before and after changes creating or removing a mismatch opposite the candidate adenosine (Fig. 3c). Nearby G*·*C-to-C*·*G substitutions 3–8 nt from the scored adenosine preserved pairing and served as controls. Uncertainty is shown as 95% bootstrap intervals over duplex families.

### Design agent

#### Design task and search budget

Reference construct: a 346-nt SINE B2 duplex from Uzonyi et al. [27], comprising a fixed scoring arm (arm 1, 146 nt), a 46-nt loop with an 8-nt barcode, a redesignable partner arm (arm 2, 146 nt), and an 8-nt tail. Target: arm 1 position 70 (0-indexed), an adenosine in a 5’-GAG-3’ context.

The agent (claude-sonnet-5-5) receives a design brief containing both reference arm sequences, the target position, construct geometry, an edit language of substitutions, insertions and deletions in original arm-2 coordinates, and constraints: arm 2 must be 106–184 nt, at most 40 edited bases, at most 20 operations, insertions interior only. The agent writes whole partner arms (up to ten per call), reads each design’s predicted change at the target, and iterates until a budget of 1,000 distinct designs is spent. Repeated designs are free (Fig. 5a).

#### Agent tools and feedback

Three tools were available. `check`: validates a design and returns the assembled sequence (no budget cost). `score`: takes up to ten designs, returns each one’s predicted editing change at the target and whether the target is paired (one budget unit per new design). `submit_final`: ends the search and returns the ten highest-scoring designs plus ten the agent never scored. The agent received a predicted change and a Boolean pairing indicator, not a structure; no folding tool was provided.

#### Control search strategies

A unit edit is one substitution, insertion, or deletion, with type and site drawn uniformly among legal choices. Four control arms use the same 1,000-design budget and ModFormer scoring function. No-tool (blind): same model and brief, never shown a score; proposes 1,000 designs that ModFormer ranks afterwards, separating design ability from feedback use. Hill climbing: starts at the reference arm, proposes *k* unit edits from its current state (*k* uniform on 1–3), accepts only a strictly higher score. Random edits: *k* unit edits from the reference arm (*k* uniform on 1–32), each draw independent. Random sequences: arm length drawn from the agent’s observed distribution, each base uniform.

#### Comparison with measured library sequences

Edit distances to measured constructs are exact unit-cost Levenshtein distances on the barcode-masked full construct, against every screened construct on the same target in the reporter library of Uzonyi et al. [27]: 1,303 of its 1,743 rows pass an exact first-arm match against the reference construct with the barcode-masked loop and tail held equal, giving 1,241 unique sequences.

### Statistical analysis

All 95% confidence intervals are percentile bootstrap, with resampling units chosen to preserve dependence: transcripts, genes, duplex families, duplexes, or constructs, as specified in the corresponding figure legends and Supplementary Table 3. Between-method comparisons used paired resampling. Intervals are pointwise and were not adjusted for multiple comparisons.

Each model was evaluated from a single fitted checkpoint (released-model seed 1313); bootstrap intervals therefore quantify sampling variation conditional on that checkpoint and do not capture between-seed training variability. The mouse-brain direct-RNA data originated from one animal, and uncertainty from site-level resampling does not reflect biological-replicate variability.

### Software

Model training ran on four NVIDIA H200 GPUs (Berzelius supercomputer, National Supercomputer Centre, Linköping University). Data processing and embedding precomputation used NVIDIA A100 GPUs on the same system. Training used Python 3.11, PyTorch 2.11.0 (CUDA 12.8), `multimolecule` 0.2.0, transformers 5.12.0, NumPy 2.4.1, SciPy 1.17.0, scikit-learn 1.8.0, pandas 3.0.0, and zarr 3.1.5, with ViennaRNA 2.7.2 for folding. The design agent is claude-sonnet-5-5.

## Data availability

Dataset sources, their roles in training and evaluation, and applicable accession identifiers are listed in Supplementary Table 1. Human and mouse RNA editing annotations were obtained from REDIportal [24]. The human Alu and invertebrate duplex datasets were obtained from the public release of Rosenwasser et al. [21]. Synthetic duplex mutational-screen data were obtained from Uzonyi et al. [27] (BioProject PRJNA649938; GEO GSE155490 for the reporter library) and Zambrano-Mila et al. [28] (BioProject PRJNA943413). Endogenous ADAR1-substrate mutational scans of AJUBA, TTYH2 and NEIL1 were obtained from Liu et al. [29] (GEO GSE138860). Benchmark datasets from published predictors were obtained from the corresponding cited studies and processed using the inclusion criteria described in Methods and Supplementary Table 3. The HEK293FT, Neuro-2a, and mouse-brain Oxford Nanopore direct-RNA sequencing data generated in this study are available in the Gene Expression Omnibus under accession GSE355083. Per-tile A-to-I labels for the direct-RNA datasets, train, validation and test split definitions for all training and evaluation sets, and source data for Figures 2–4 and Extended Data Figures 1–3 are available at Zenodo (doi:10.5281/zenodo.24300820) under CC BY-NC 4.0.

## Code availability

ModFormer model weights, the released model checkpoint and the jointly fine-tuned checkpoint (both seed 1313), are available at Zenodo (doi:10.5281/zenodo.24300574) under the Creative Commons Attribution-NonCommercial 4.0 International license (CC BY-NC 4.0). The checkpoint excludes the frozen RiNALMo-giga backbone weights, which are obtained from their original release through `multimolecule` 0.2.0. Code to recreate the figures and to use ModFormer for custom predictions is available at https://github.com/nygctech/ModFormer under the MIT license.

## Acknowledgments

This work was supported by the Knut and Alice Wallenberg Foundation, Swedish Society for Medical Research, Swedish Research Council, Science for Life Laboratory, and the MacMillan Center for the Non-Coding Genome. Computational resources were provided on the Berzelius system funded by the Knut and Alice Wallenberg Foundation and operated by NAISS. S.V. is additionally supported as a Wallenberg Academy Fellow and SciLifeLab Fellow at Uppsala University.

## Author contributions

C.S. and S.V. conceptualized the project; C.S. designed and developed ModFormer with contributions from E.N., B.Y., N.D.B., M.G. and supervision from A.D. and S.V.; B.Y. contributed to the windowed transcript embedding; N.D.B. contributed to the sequence-perturbation analysis of editing logits; E.N. contributed to model testing and evaluation; M.G. contributed to data processing; C.S. and L.E. performed direct RNA sequencing experiments; C.S. conducted all other experiments and analyses; J.H.L. performed wet-lab experiments and reviewed the manuscript; C.S., D.D. and S.V. coordinated the project; C.S., A.D., E.N., and S.V. wrote the manuscript; all authors discussed the results.

## Additional information

### Supplementary information

**Supplementary Table 1**. Datasets used for ModFormer training, adaptation, and evaluation.

**Supplementary Table 2**. ModFormer architecture, training, and adaptation settings.

**Supplementary Table 3**. Published-model benchmarks, prediction settings, and statistical comparisons.

## Extended Data Figures

**Extended Data Fig. 1.**
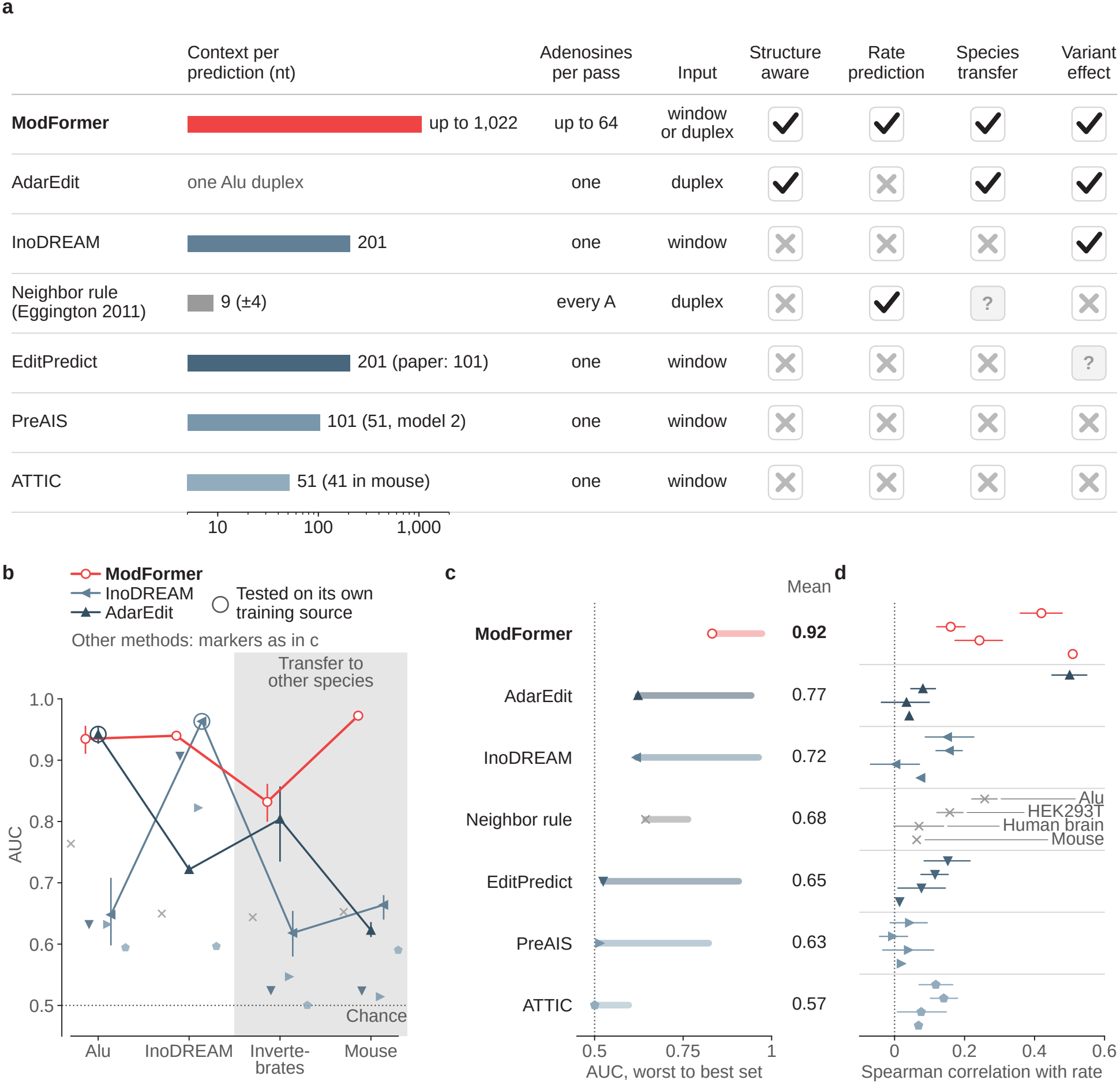
ModFormer performance against published editing-site predictors. **a**, Inputs and reported capabilities of ModFormer and six comparators: AdarEdit [21], InoDREAM [33], ATTIC [34], EditPredict [35], PreAIS [36], and a neighbor rule [11]. Bars, input context (nt; log scale); checkmarks, shown in the paper; crosses, not shown; question marks, claimed without results or compared only qualitatively. AdarEdit reports human-to-invertebrate transfer but primarily trains species-specific models. **b**, Editing-site AUC across Alu (729 sites), InoDREAM (17,956), invertebrate (1,678), and mouse Neuro-2a/brain (21,667) benchmarks. Gray, species absent from model training; rings, evaluation on a method’s training-data source; dotted line, chance. Markers: ModFormer, open circle; InoDREAM, left-pointing triangle; AdarEdit, upward triangle; neighbor rule, cross; EditPredict, downward triangle; PreAIS, right-pointing triangle; ATTIC, pentagon. ModFormer, AdarEdit, and InoDREAM have 95% bootstrap intervals. Cohorts and labels differ from Fig. 2a. **c**, Minimum, maximum, and unweighted mean AUC across benchmarks; methods ordered by mean. **d**, Spearman *ρ* with measured editing rates at edited sites in Alu (1,683), HEK293T (2,461), human brain (775), and mouse Neuro-2a/brain (276,924). Each block has four sub-rows, top to bottom: Alu, HEK293T, human brain, mouse (named once, on the neighbor-rule block).

**Extended Data Fig. 2.**
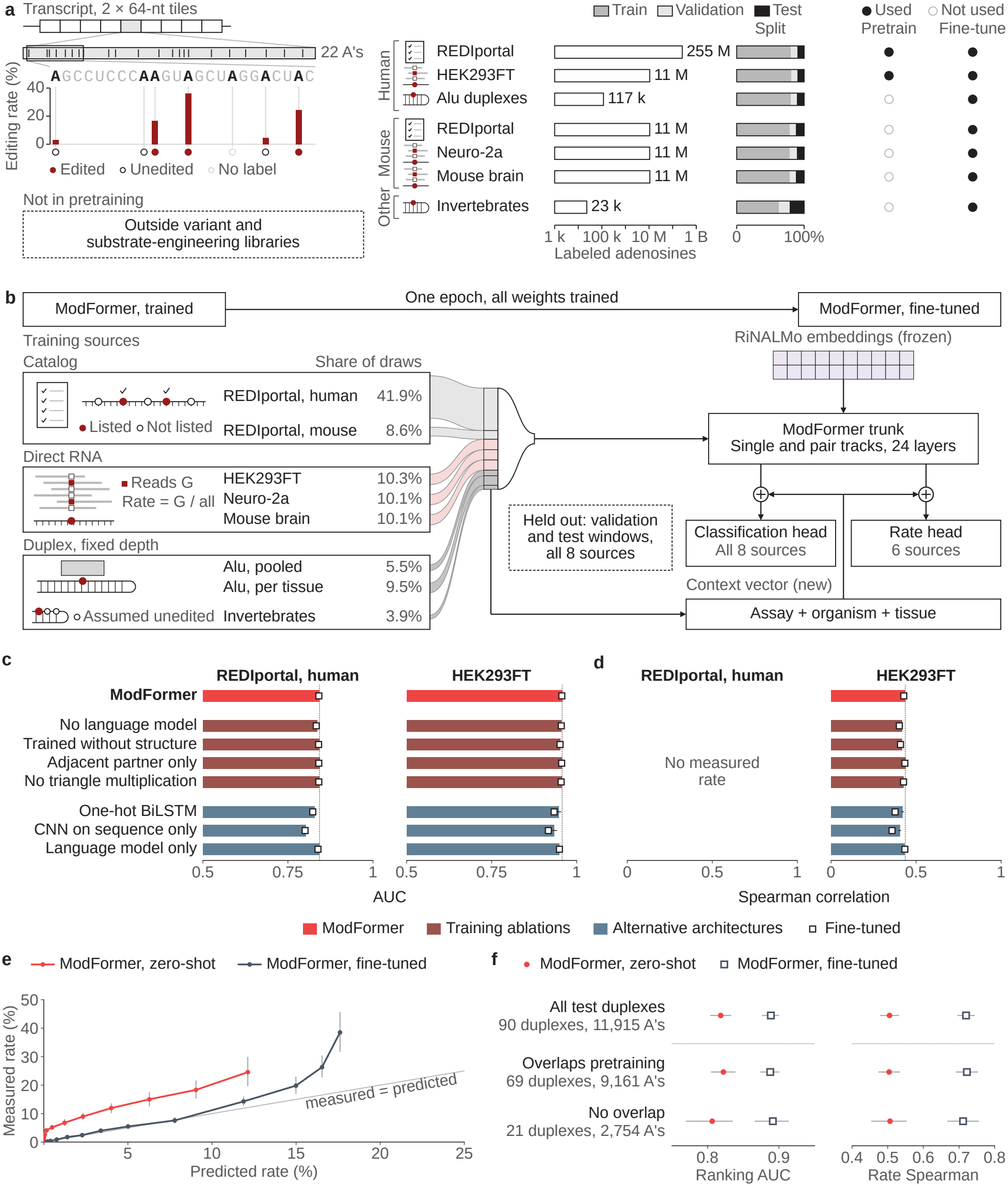
Training data, fine-tuning, and held-out editing predictions. Uncertainty displays are 95% percentile-bootstrap intervals (thin lines). **a**, Datasets, label definitions, sizes, and train/validation/test splits. Anchor sites are scored alongside their predicted 64-nt pairing partner; filled, hollow, and dotted adenosines are edited, unedited, and unlabeled. Filled circles mark training uses. Dashed box, external variant and substrate-engineering datasets [6, 21, 27–29], excluded from pretraining; Rosenwasser Alu and invertebrate duplexes contribute to fine-tuning. REDIportal tissues follow D’Addabbo et al. [24]. **b**, One-epoch joint fine-tuning over eight sources; band height, sampling share. Assay/context vectors enter both prediction heads; held-out splits are excluded. **c**,**d**, Editing-site AUC (**c**) and rate Spearman *ρ* (**d**) on held-out human REDIportal and HEK293FT datasets. Bars, zero-shot; squares, fine-tuned; dotted lines, ModFormer’s zero-shot value. REDIportal lacks measured rates. **e**, Mean predicted versus measured editing rates for 11,915 adenosines in 90 Alu duplexes, each point is a bin mean over deciles then the top 1%; diagonal, equal means. Group means do not measure per-site accuracy. **f**, Ranking AUC and rate *ρ* in all 90 Alu duplexes, 69 sharing a 128-nt window with the pretraining tiles and 21 without. Red dot, zero-shot; open square, one-epoch fine-tune; no line joins them. Without versus with overlap (zero-shot): AUC, 0.806 versus 0.822; *ρ*, 0.506 versus 0.504.

**Extended Data Fig. 3.**
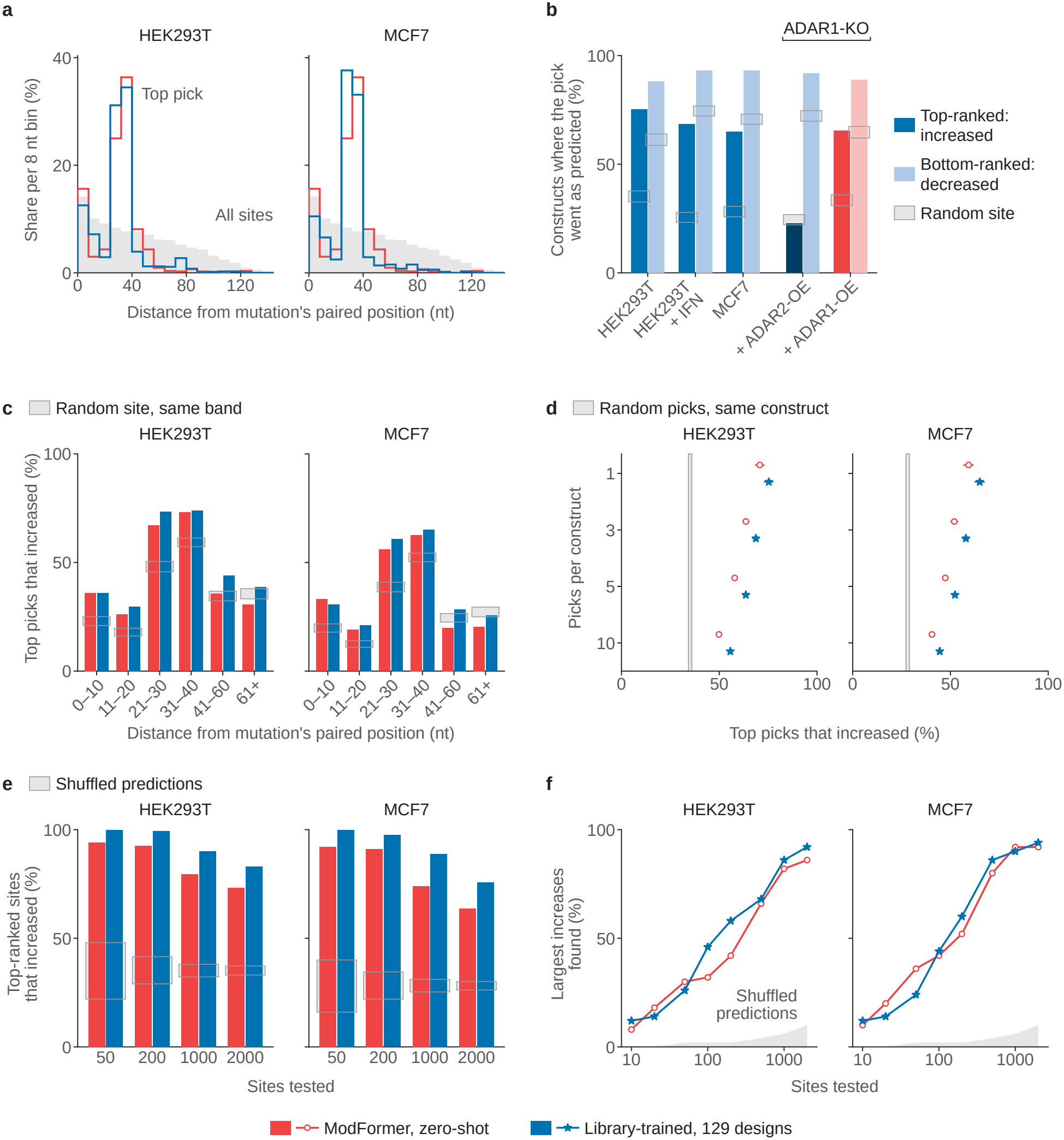
Editing-site prioritization across mutation distances and cellular contexts. Held-out library constructs; red, zero-shot ModFormer; blue, library-trained model. Top and bottom picks are the sites with greatest predicted editing gain and loss. **a**, Distance from selected sites to the position paired with the nearest mutation (8-nt bins); gray, all candidate sites. **b**, Fraction of constructs with observed editing changes in the predicted direction across five cellular conditions. Dark/light, top/bottom picks; gray, random baseline. The first four conditions use library-trained predictions; the dark navy bar is the library-trained top pick in ADAR1 knockout with ADAR2 overexpression. The red/pink pair (ADAR1 knockout with ADAR1 overexpression) uses ModFormer zero-shot predictions only, as the library-trained model has no ADAR1-overexpression arm, so enzyme contexts are not compared using identical checkpoints. **c**, Fraction of top picks showing increased editing with one pick per mutation-distance bin; gray, distance-matched random picks. **d**, Fraction showing increased editing versus number of picks per construct; 95% bootstrap intervals and matched random baseline. **e**, Fraction of ranked sites with increased editing versus total sites tested; gray, shuffled rankings. **f**, Recovery of the 50 largest measured gains versus sites tested; gray, shuffled rankings. Panels **a**,**c–f** show HEK293T and MCF7. Within 10 nt of the mutation-paired position, the difference in top-pick gain fraction (library-trained minus zero-shot; 95% intervals) was *−*0.001 (*−*0.027 to +0.025) in HEK293T, +0.018 (*−*0.007 to +0.044) with interferon, *−*0.024 (*−*0.050 to +0.003) in MCF7, and +0.030 (+0.001 to +0.060) with ADAR2 overexpression.

**Supplementary Table 1.**
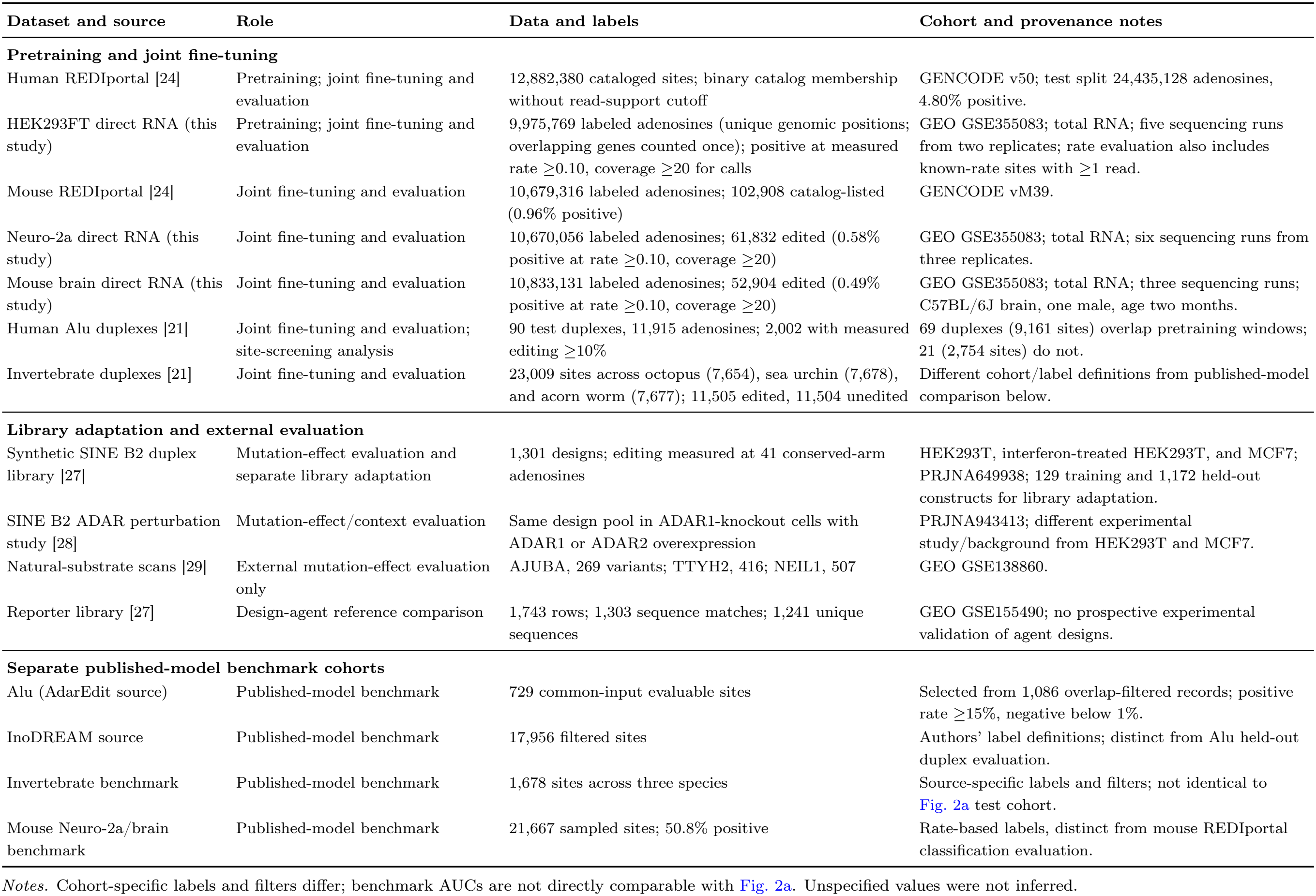
Datasets used for ModFormer training, adaptation, and evaluation.

**Supplementary Table 2.**
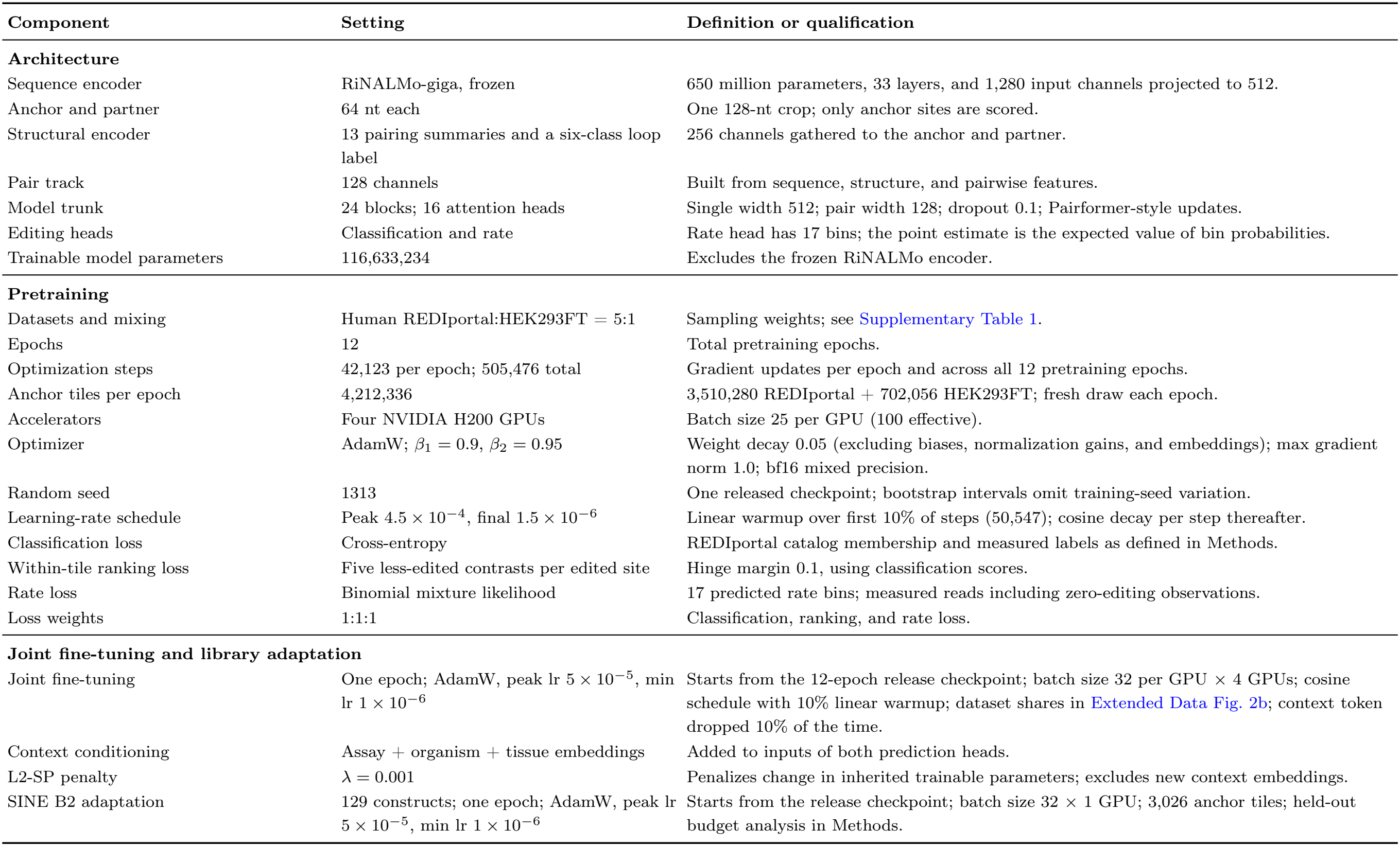
ModFormer architecture, training, and adaptation settings.

**Supplementary Table 3.**
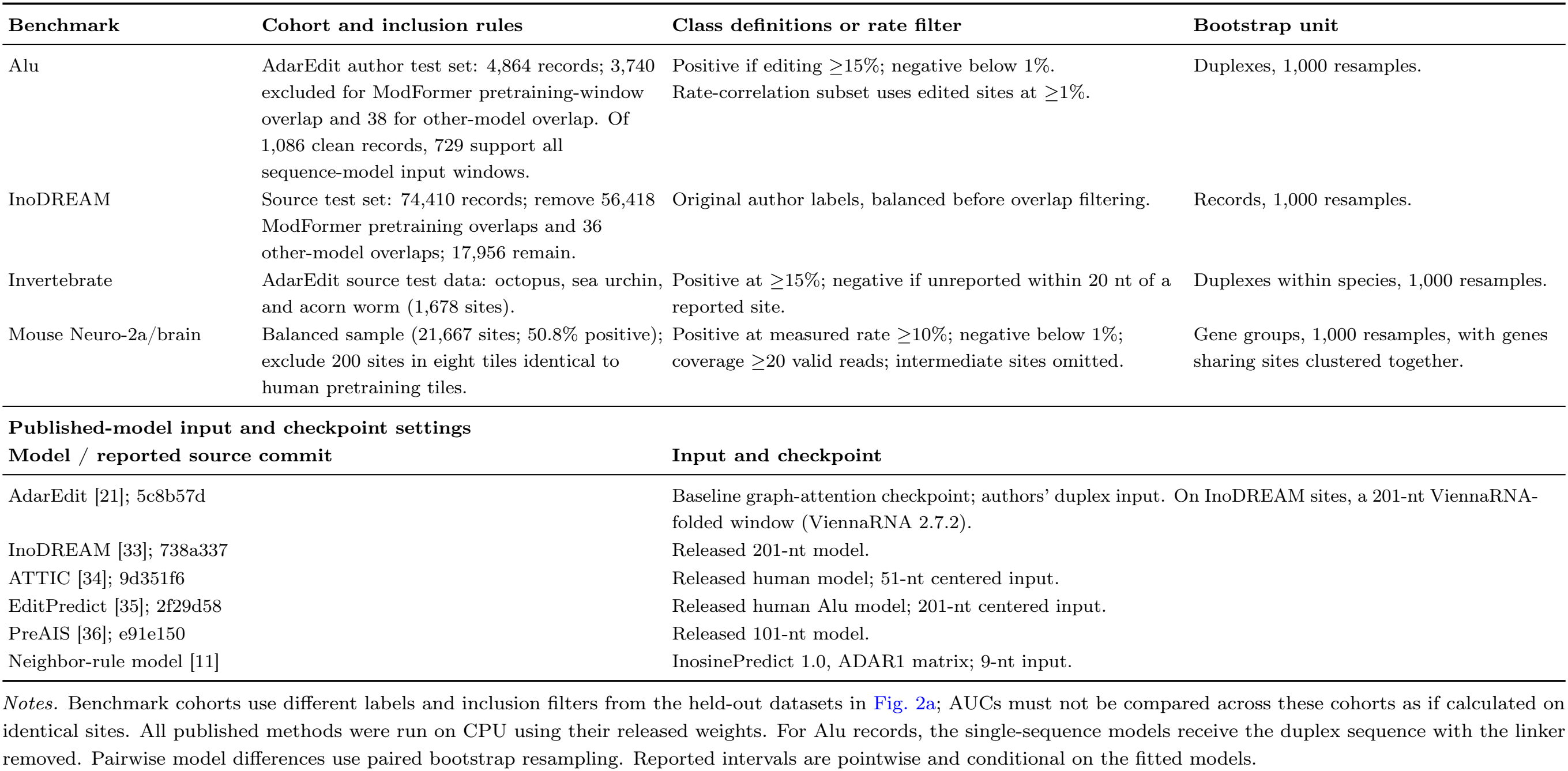
Published-model benchmarks, prediction settings, and statistical comparisons.

