## Supplemental Tables for "ModFormer: A unified structure-aware model of transcriptome-wide A-to-I RNA editing"

### Supplementary Tables

Supplementary Table 1: Datasets used for ModFormer training, adaptation, and evaluation.

| Dataset and source | Role | Data and labels | Cohort and provenance notes |
|---|---|---|---|
| **Pretraining and joint fine-tuning** | | | |
| Human REDIportal [1] | Pretraining; joint fine-tuning and evaluation | 12,882,380 cataloged sites; binary catalog membership without read-support cutoff | GENCODE v50; test split 24,435,128 adenosines, 4.80% positive. |
| HEK293FT direct RNA (this study) | Pretraining; joint fine-tuning and evaluation | 9,975,769 labeled adenosines (unique genomic positions; overlapping genes counted once); positive at measured rate ≥0.10, coverage ≥20 for calls | GEO GSE355083; total RNA; five sequencing runs from two replicates; rate evaluation also includes known-rate sites with ≥1 read. |
| Mouse REDIportal [1] | Joint fine-tuning and evaluation | 10,679,316 labeled adenosines; 102,908 catalog-listed (0.96% positive) | GENCODE vM39. |
| Neuro-2a direct RNA (this study) | Joint fine-tuning and evaluation | 10,670,056 labeled adenosines; 61,832 edited (0.58% positive at rate ≥0.10, coverage ≥20) | GEO GSE355083; total RNA; six sequencing runs from three replicates. |
| Mouse brain direct RNA (this study) | Joint fine-tuning and evaluation | 10,833,131 labeled adenosines; 52,904 edited (0.49% positive at rate ≥0.10, coverage ≥20) | GEO GSE355083; total RNA; three sequencing runs; C57BL/6J brain, one male, age two months. |
| Human Alu duplexes [2] | Joint fine-tuning and evaluation; site-screening analysis | 90 test duplexes, 11,915 adenosines; 2,002 with measured editing ≥10% | 69 duplexes (9,161 sites) overlap pretraining windows; 21 (2,754 sites) do not. |
| Invertebrate duplexes [2] | Joint fine-tuning and evaluation | 23,009 sites across octopus (7,654), sea urchin (7,678), and acorn worm (7,677); 11,505 edited, 11,504 unedited | Different cohort/label definitions from published-model comparison below. |
| **Library adaptation and external evaluation** | | | |
| Synthetic SINE B2 duplex library [3] | Mutation-effect evaluation and separate library adaptation | 1,301 designs; editing measured at 41 conserved-arm adenosines | HEK293T, interferon-treated HEK293T, and MCF7; PRJNA649938; 129 training and 1,172 held-out constructs for library adaptation. |
| SINE B2 ADAR perturbation study [4] | Mutation-effect/context evaluation | Same design pool in ADAR1-knockout cells with ADAR1 or ADAR2 overexpression | PRJNA943413; different experimental study/background from HEK293T and MCF7. |
| Natural-substrate scans [5] | External mutation-effect evaluation only | AJUBA, 269 variants; TTYH2, 416; NEIL1, 507 | GEO GSE138860. |
| Reporter library [3] | Design-agent reference comparison | 1,743 rows; 1,303 sequence matches; 1,241 unique sequences | GEO GSE155490; no prospective experimental validation of agent designs. |
| **Separate published-model benchmark cohorts** | | | |
| Alu (AdarEdit source) | Published-model benchmark | 729 common-input evaluable sites | Selected from 1,086 overlap-filtered records; positive rate ≥15%, negative below 1%. |
| InoDREAM source | Published-model benchmark | 17,956 filtered sites | Authors' label definitions; distinct from Alu held-out duplex evaluation. |
| Invertebrate benchmark | Published-model benchmark | 1,678 sites across three species | Source-specific labels and filters; not identical to Fig. a test cohort. |
| Mouse Neuro-2a/brain benchmark | Published-model benchmark | 21,667 sampled sites; 50.8% positive | Rate-based labels, distinct from mouse REDIportal classification evaluation. |

*Notes.* Cohort-specific labels and filters differ; benchmark AUCs are not directly comparable with Fig. a. Unspecified values were not inferred.

Supplementary Table 2: ModFormer architecture, training, and adaptation settings.

| **Component** | **Setting** | **Definition or qualification** |
|---|---|---|
| **Architecture** | | |
| Sequence encoder | RiNALMo-giga, frozen | 650 million parameters, 33 layers, and 1,280 input channels projected to 512. |
| Anchor and partner | 64 nt each | One 128-nt crop; only anchor sites are scored. |
| Structural encoder | 13 pairing summaries and a six-class loop label | 256 channels gathered to the anchor and partner. |
| Pair track | 128 channels | Built from sequence, structure, and pairwise features. |
| Model trunk | 24 blocks; 16 attention heads | Single width 512; pair width 128; dropout 0.1; Pairformer-style updates. |
| Editing heads | Classification and rate | Rate head has 17 bins; the point estimate is the expected value of bin probabilities. |
| Trainable model parameters | 116,633,234 | Excludes the frozen RiNALMo encoder. |
| **Pretraining** | | |
| Datasets and mixing | Human REDIportal:HEK293FT = 5:1 | Sampling weights; see Supplementary Table. |
| Epochs | 12 | Total pretraining epochs. |
| Optimization steps | 42,123 per epoch; 505,476 total | Gradient updates per epoch and across all 12 pretraining epochs. |
| Anchor tiles per epoch | 4,212,336 | 3,510,280 REDIportal + 702,056 HEK293FT; fresh draw each epoch. |
| Accelerators | Four NVIDIA H200 GPUs | Batch size 25 per GPU (100 effective). |
| Optimizer | AdamW; $\beta_1 = 0.9$, $\beta_2 = 0.95$ | Weight decay 0.05 (excluding biases, normalization gains, and embeddings); max gradient norm 1.0; bf16 mixed precision. |
| Random seed | 1313 | One released checkpoint; bootstrap intervals omit training-seed variation. |
| Learning-rate schedule | Peak $4.5 \times 10^{-4}$, final $1.5 \times 10^{-6}$ | Linear warmup over first 10% of steps (50,547); cosine decay per step thereafter. |
| Classification loss | Cross-entropy | REDIportal catalog membership and measured labels as defined in Methods. |
| Within-tile ranking loss | Five less-edited contrasts per edited site | Hinge margin 0.1, using classification scores. |
| Rate loss | Binomial mixture likelihood | 17 predicted rate bins; measured reads including zero-editing observations. |
| Loss weights | 1:1:1 | Classification, ranking, and rate loss. |
| **Joint fine-tuning and library adaptation** | | |
| Joint fine-tuning | One epoch; AdamW, peak lr $5 \times 10^{-5}$, min lr $1 \times 10^{-6}$ | Starts from the 12-epoch release checkpoint; batch size 32 per GPU $\times$ 4 GPUs; cosine schedule with 10% linear warmup; dataset shares in Extended Data Fig. b; context token dropped 10% of the time. |
| Context conditioning | Assay + organism + tissue embeddings | Added to inputs of both prediction heads. |
| L2-SP penalty | $\lambda = 0.001$ | Penalizes change in inherited trainable parameters; excludes new context embeddings. |
| SINE B2 adaptation | 129 constructs; one epoch; AdamW, peak lr $5 \times 10^{-5}$, min lr $1 \times 10^{-6}$ | Starts from the release checkpoint; batch size 32 $\times$ 1 GPU; 3,026 anchor tiles; held-out budget analysis in Methods. |

Supplementary Table 3: Published-model benchmarks, prediction settings, and statistical comparisons.

| **Benchmark** | **Cohort and inclusion rules** | **Class definitions or rate filter** | **Bootstrap unit** |
|---|---|---|---|
| Alu | AdarEdit author test set: 4,864 records; 3,740 excluded for ModFormer pretraining-window overlap and 38 for other-model overlap. Of 1,086 clean records, 729 support all sequence-model input windows. | Positive if editing $\geq$15%; negative below 1%. Rate-correlation subset uses edited sites at $\geq$1%. | Duplexes, 1,000 resamples. |
| InoDREAM | Source test set: 74,410 records; remove 56,418 ModFormer pretraining overlaps and 36 other-model overlaps; 17,956 remain. | Original author labels, balanced before overlap filtering. | Records, 1,000 resamples. |
| Invertebrate | AdarEdit source test data: octopus, sea urchin, and acorn worm (1,678 sites). | Positive at $\geq$15%; negative if unreported within 20 nt of a reported site. | Duplexes within species, 1,000 resamples. |
| Mouse Neuro-2a/brain | Balanced sample (21,667 sites; 50.8% positive); exclude 200 sites in eight tiles identical to human pretraining tiles. | Positive at measured rate $\geq$10%; negative below 1%; coverage $\geq$20 valid reads; intermediate sites omitted. | Gene groups, 1,000 resamples, with genes sharing sites clustered together. |

**Published-model input and checkpoint settings**

| **Model / reported source commit** | **Input and checkpoint** |
|---|---|
| AdarEdit [2]; 5c8b57d | Baseline graph-attention checkpoint; authors' duplex input. On InoDREAM sites, a 201-nt ViennaRNA-folded window (ViennaRNA 2.7.2). |
| InoDREAM [6]; 738a337 | Released 201-nt model. |
| ATTIC [7]; 9d351f6 | Released human model; 51-nt centered input. |
| EditPredict [8]; 2f29d58 | Released human Alu model; 201-nt centered input. |
| PreAIS [9]; e91e150 | Released 101-nt model. |
| Neighbor-rule model [10] | InosinePredict 1.0, ADAR1 matrix; 9-nt input. |

*Notes.* Benchmark cohorts use different labels and inclusion filters from the held-out datasets in Fig. a; AUCs must not be compared across these cohorts as if calculated on identical sites. All published methods were run on CPU using their released weights. For Alu records, the single-sequence models receive the duplex sequence with the linker removed. Pairwise model differences use paired bootstrap resampling. Reported intervals are pointwise and conditional on the fitted models.
